# Immediate impact of the 2021 harmful algal bloom in southeast Hokkaido on the rocky intertidal community and its spatial variation

**DOI:** 10.1101/2023.12.01.569513

**Authors:** Yuan Yao, Takashi Noda

**Author notes:** **Corresponding author:** Given name: Yuan, Family name: Yao, E-mail address, Affiliation address: Graduate School of Environmental Science, Hokkaido University, N10W5, Kita-ku, Sapporo, Hokkaido 060-0810, Japan.

## Abstract

There has been a limited number of studies on the effects of harmful algal blooms (HABs) on natural rocky intertidal ecosystems. From mid-September to early November 2021, an unprecedented HAB caused by *Karenia selliformis* hit the Pacific coast of southeast Hokkaido, Japan, for the first time, causing massive mortalities among marine organisms. To clarify the immediate impacts of the HAB on abundance of 10 rocky intertidal species in four functional groups (macroalgae, sessile invertebrates, molluscan grazers, and molluscan carnivores), we focused on two questions. 1) How did the HAB affect the abundance of each species differently at the regional scale? 2) How did the impacts of the HAB on the abundance of each functional groups vary spatially, and was the spatial variation of the HAB impacts related to the spatial distribution of cell density of HAB species? To study these issues, we compared census data for 17 years before the HAB and within one month after it for five shores on the southeast coast of Hokkaido. Results showed that two macroalgae species and all three molluscan grazer species declined significantly after the HAB. Moreover, the decrease of molluscan grazers was significantly correlated with cell density. These results suggest that the impacts of the HAB in southeast Hokkaido on abundance of rocky intertidal organisms are highly variable depending on species and locality, presumably because of differences in species-specific tolerances to HAB toxins and spatial variation in the density of the HAB organisms.

## 1. Introduction

Harmful algal blooms (HABs) are known as pernicious disturbance events for aquatic organisms and coastal ecosystems, and they cause substantial ecological alteration and financial losses. Many previous studies focused on the negative impacts of HABs on commercial species and explored the influence of HABs on abundance, survival, and physical condition (Ohgaki *et al*., 2019; Trottet *et al*., 2021), and the underlying mechanisms behind these negative influences (Landsberg, 2002; Gobler *et al*., 2004; Spilmont *et al*., 2009). In contrast, there are only a limited number of studies examining the ecological consequences of HABs (Simon & Dauer, 1972; Wear & Gardner, 2001; Branch *et al*., 2013).

To understand the ecological consequences of HABs in coastal ecosystems, a fundamental and crucial task is to elucidate the immediate impact of HABs on the population sizes of multiple species at multiple localities at a regional scale because understanding this impact should improve our understanding of how HABs directly affect the community. The immediate impacts often do not include indirect effects on population dynamics triggered by species interactions. In addition, most coastal organisms form meta-populations consisting of many local populations in which every population experiences a locally variable environment (Sale *et al*., 2006). Therefore, knowledge about the immediate impacts from direct effects of HABs and the long-term impacts caused by subsequent species interactions can be inferred by following changes in the abundance of every species at multiple localities in a region.

From mid-September to early November 2021, there was an unprecedented large-scale HAB with *Karenia selliformis* as main species in the coastal waters off the southeast coast of Hokkaido, Japan (Iwataki *et al*., 2022; Kuroda *et al*., 2022). This was the first time such a severe HAB had been recorded in this area (Kuroda *et al*., 2021). Because this HAB caused serious fishery losses, as exemplified by those of sea urchins living in the intertidal and subtidal zone (more than 17 billion JPY), it might have also affected various organisms living in rocky intertidal habitats along southeast Hokkaido. Previous studies of the impacts of HABs on rocky intertidal communities indicate that mobile consumers, such as sea urchins and starfish, are susceptible to the effects of HABs, whereas sessile invertebrates, such as mussels and oysters, that compete with macroalgae (i.e., primary producers) for substrate space, are resilient (Crowe *et al*., 2000; Worm & Lotze, 2006; Ohgaki *et al*., 2019). However, none of these studies used long-term census data to elucidate the immediate impact of HABs on population sizes of multiple species from different functional groups at multiple sites at a regional scale.

*Karenia selliformis* may cause massive mortality in macroalgae and mollusks. The latest study of the same HAB event in southeast Hokkaido reported that *K. selliformis* has a lethal effect on juvenile kelp (*Saccharina japonica* and *S. sculpera*) caused by bleaching juvenile kelp thalli and subsequent cell death (Natsuike *et al*., 2023). This finding implies that *K. selliformis* may have a lethal effect on other macroalgae living in rocky intertidal habitats. Moreover, previous studies demonstrated that *K. selliformis* have a lethal effect on many mollusks, including limpets and snails (Vellojin *et al*., 2023), which are the dominant mollusks in rocky intertidal communities. Furthermore, a study of the same HAB event also observed intensive mortalities of limpets and whelks immediately after the HAB occurrence (Misaka and Ando, 2021). Although none of the above-mentioned studies fully explained the specific toxicological mechanism of *K. selliformis* on these species, it is reasonable to conclude that *K. selliformis* caused these massive mortalities. Because the functional groups of organisms mentioned in previous studies (i.e., macroalgae [kelp], molluscan grazers [limpets], and molluscan carnivores [snails]) are dominant in rocky intertidal communities in southeast Hokkaido, assessing the immediate changes in their abundance after this large HAB could provide important information for exploring subsequent changes in species interactions and population dynamics.

The purpose of this study was to clarify the immediate impacts of the 2021 HAB in southeast Hokkaido on different functional groups including macroalgae (i.e., primary producers), molluscan grazers (i.e., primary consumers), molluscan carnivores (i.e., secondary consumers), and sessile invertebrates (i.e., competitors with macroalgae for space) in rocky intertidal communities, focusing especially on any changes in population size among different species, functional groups, and localities. For this purpose, the following questions about the immediate impacts of the HAB at the population level were addressed. 1) How did the HAB affect the population size of each species at a regional scale and were there differences among functional groups? 2) How did the impacts of the HAB on the abundance of each functional group vary depending on shore, and were spatial variations of the HAB impacts related to the spatial distribution of cell density of HAB species? To answer these questions, long-term census data from southeast Hokkaido for 17 years before the HAB (from 2004 to 2020) and sample data from within one month after HAB were analyzed. These large-scale long-term monitoring census data enable estimation of the changes of population sizes, in which differences in population size between years with and without the influence of the HAB are standardized to population variability under normal conditions for every species at each location (Iwasaki *et al*., 2016; Iwasaki & Noda, 2018).

## 2. Material and methods

### 2.1 Study area

Our study sites were located along over 50 km of shoreline on the Pacific coast of eastern Hokkaido (Fig. 1), within the range of the HAB in September 2021. At these study sites, there are various rocky intertidal organisms, including macroalgae, sessile invertebrates (such as barnacles), molluscan grazers, and molluscan carnivores (Nakaoka *et al*., 2006). These species are distributed in the mid-tidal zone within a vertical range of tens of centimeters. The horizontal extent of our study sites should have covered the entire range of the metapopulation of most macroalgae and benthic mollusks (Kinlan and Gaines, 2003).

**Fig. 1.**
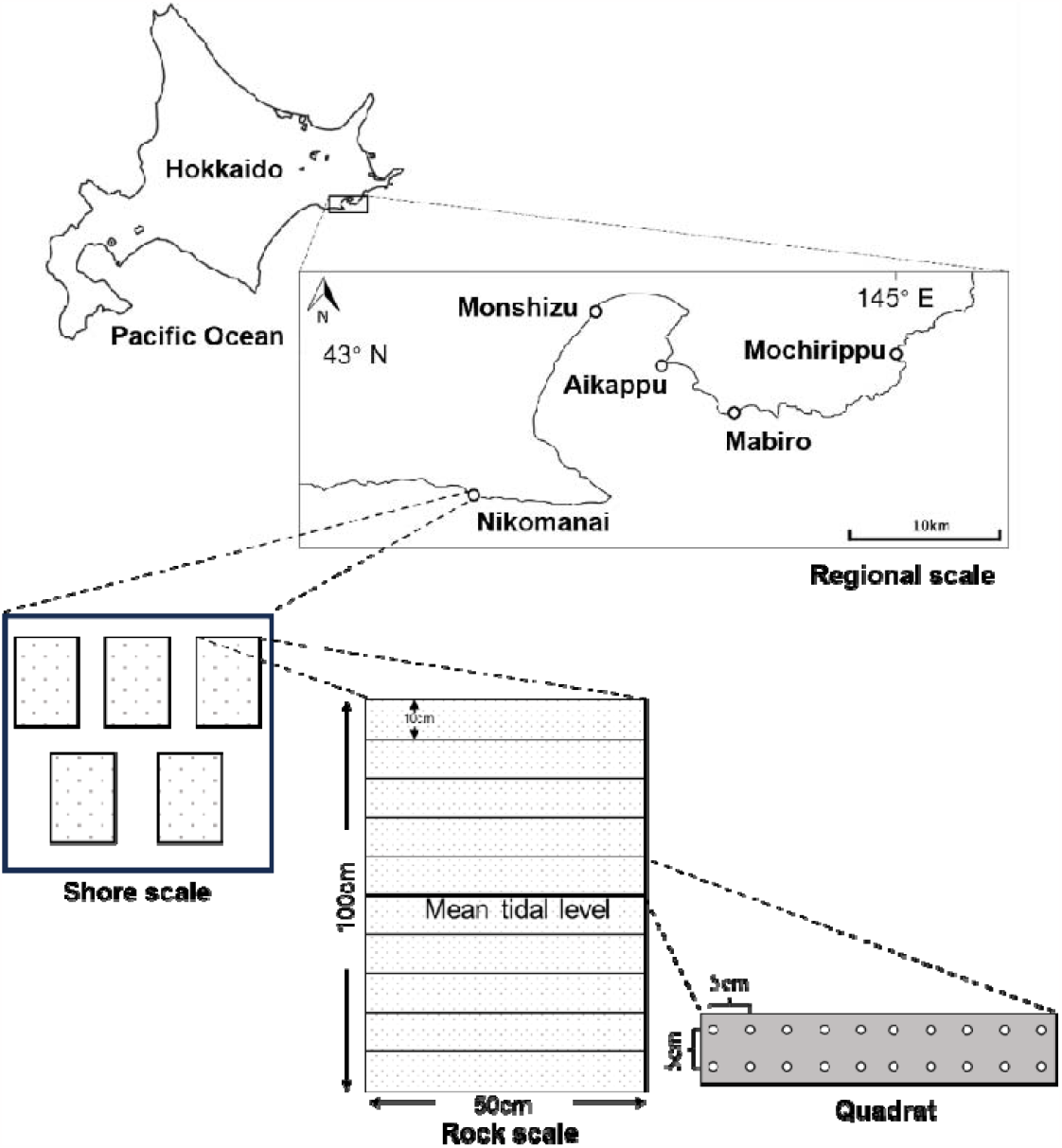
Diagram of study site locations and census design. Five shores were chosen along the Pacific coast of eastern Hokkaido, Japan. At each shore, five permanent plots (50 cm wide by 100 cm high) were set up on nearly vertical rocks (25 rocks in total), with the vertical midpoints corresponding to the mean tidal level. Each plot was evenly divided vertically into 10 quadrats (50 cm wide by 10 cm high). The white circles in the quadrat diagram represent fixed observation points.

### 2.2 Census design

Prior to the 2021 HAB, sampling following a hierarchical design (Noda, 2004) was conducted at five shores on the Pacific coast of southeast Hokkaido (Mochirippu [MC]: 43°01′02′′N, 145°01′30′′E; Mabiro [MB]: 42°59′14′′N, 144°53′24′′E; Aikappu [AP]: 43°01′02′′N, 144°50′02′′E; Monshizu [MZ]: 43°02′59′′N, 144°46′42′′ E; and Nikomanai [NN]: 42°56′23′′N, 144°40′26′′E) (Fig. 1) beginning in 2004. At each shore, 5 permanent plots 50 cm wide by 100 cm high were set up on nearly vertical rocks (25 plots in total), with the vertical midpoints corresponding to the mean tidal level. The distance between neighboring plots ranged from 6.6 to 113 m (mean ± SD, 34.1 m ± 27.8 m). Each plot was evenly divided vertically into 10 quadrats (50 cm wide by 10 cm high). For all censuses, the number of individuals of each mobile species (i.e., molluscan grazers and molluscan carnivores) was obtained for each quadrat, representing the vertical sections of each permanent plot at the rock scale (Fig. 1). In addition, the coverage of each sessile species (i.e., macroalgae and sessile invertebrates) was estimated by using a point-sampling method in which each quadrat in a plot was divided into 20 fixed observation points with 2 horizontal rows and 10 vertical columns. The interval between each point was 5 cm both vertically and horizontally. The dominant species occupying each fixed observation point was recorded. Therefore, the coverage of each species was the product of the number of recorded points and25 cm^2^, and the coverage was used as the abundance of sessile species. The abundance of rocky intertidal organisms in permanent plots were determined annually at low tide in late October or early November. In November 2021, within 1 month after the HAB off the Pacific coast of eastern Hokkaido, the same census was performed at these sites to assess the immediate impact of the HAB.

### 2.3 Species selection

To avoid inaccurate assessment of the effect of the HAB on population size, only the species at the regional scale and functional groups at the shore scale that were relatively abundant and observed continuously were selected as the subject species. For sessile species, a threshold coverage of 125 cm^2^ at the regional scale in each year was specified, and for mobile species, the threshold was 5 individuals. According to past census data in this area, the sessile invertebrate *Chthamalus dalli*; macroalgae *Corallina pilulifera, Chondrus yendoi, Gloiopeltis furcata, Heterochordaria abietina*, and *Polysiphonia yendoi*; molluscan grazers *Littorina sitkana, Lottia cassis*, and *Lottia* sp. (the scientific name for this limpet has not been settled; Yamazaki, 2011); and the molluscan carnivore *Nucella lima* meet the threshold values at the regional scale. In addition, the abundance of each functional group was recorded as the total abundance of all species within that functional group at the shore scale. To ensure the accuracy of the assessment, we excluded functional groups that have not been observed for two or more consecutive years. According to past census data in this area, sessile invertebrates, macroalgae, and molluscan grazers meet these threshold values at the shore scale.

### 2.4 Calculation of cell density of HAB species

The cell density of HAB species was derived by using the surface chlorophyll-a concentrations reported by Kuroda *et al*. (2022), who measured chlorophyll-*a* concentration along a ship track (during 4–14 October 2021) in their study of the same HAB event. They presented chlorophyll-*a* concentration values in a census map as colored grid cells (Fig. 2A in the present study, Figure 4 in Kuroda *et al*., 2022.). In addition, they provided a log-log regression equation of *Karenia* spp. abundance versus chlorophyll-*a* concentration (Figure 7 in Kuroda *et al*., 2022):

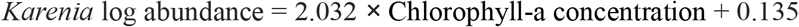

In the present study, we selected the colored grid cell perpendicular to the coastline of each shore to calculate the cell density of *Karenia* spp. Specifically, we first calculated the distance between the RGB color space of a selected colored grid cell and the RGB color space of each colored grid cell in the legend. Then, the integer of the colored grid cell in the legend with the minimum distance from the selected colored grid cell was used as the chlorophyll-a concentration (Fig. 2B). Finally, the log abundance of *Karenia* spp. was calculated by the above regression equation, and we conducted an exponential operation to calculate the cell density of *Karenia* spp.

**Fig. 2.**
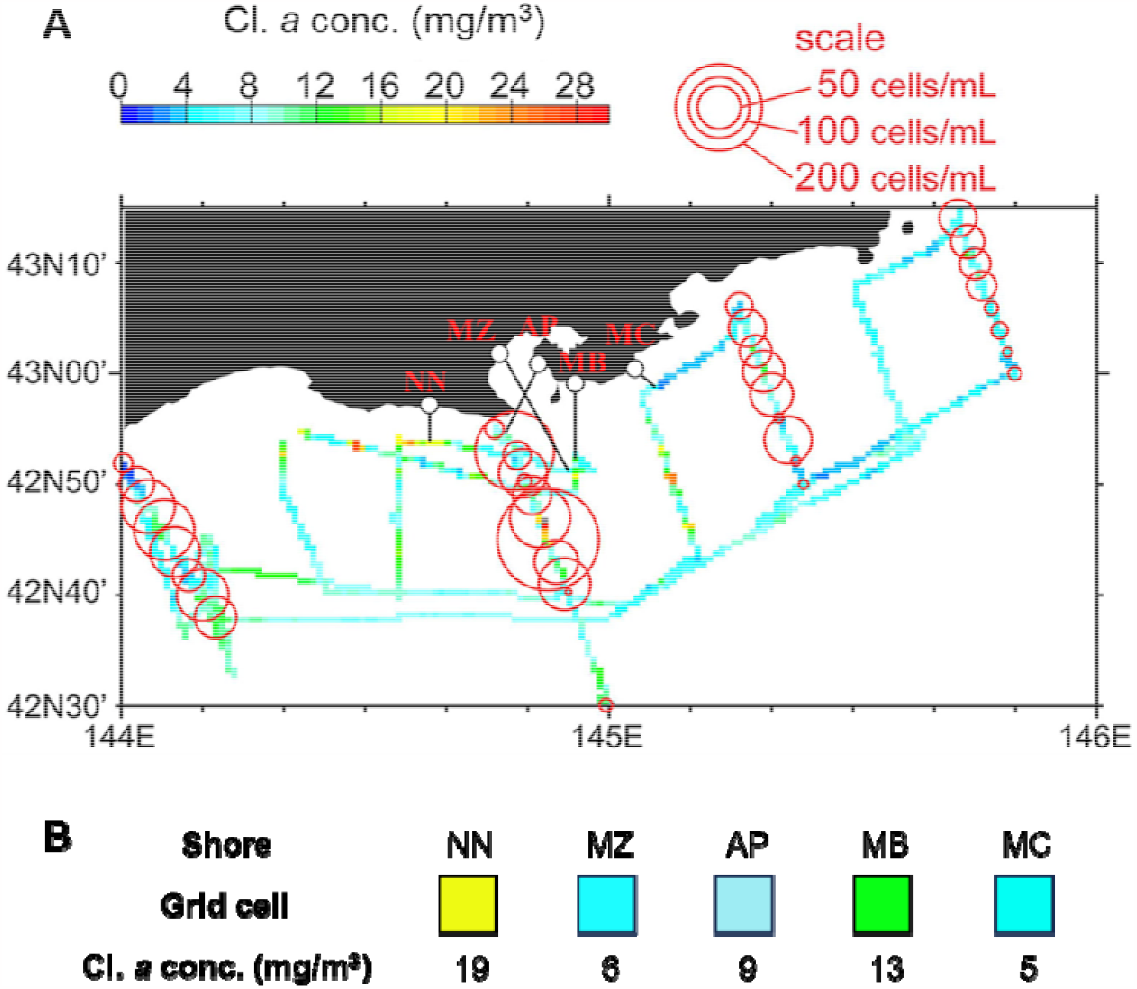
(color should be used in print). (A) Surface chlorophyll-a concentrations (colored grid cells) along the ship track during 4–14 October, 2021 conducted by Kuroda et al. Grid cells shows averages across 45″ (longitude) × 30″ (latitude) (i.e., an approximately 1 × 1 km rectangle). Karenia spp. abundance at a depth of 10 m is also shown by the area of red circles. We added five black hollow circles in original figure to indicate the census shore in present study. And we added the black solid lines perpendicular to the shoreline of census shores to represent the colored grid cells that used to calculate the cell density of Karenia spp. in present study. (B) The enlarged view of the color grid cells, and the chlorophyll a concentration calculated by RGB color value.

### 2.5 Statistical analysis

To evaluate the effects of HAB on each species (*i*) at the regional scale (*ES*_*REGION,i*_) and each functional group (*j*) at each shore (*k*) (*ES*_*SHORE,j,k*_), we calculated the effect sizes (*ES*) of the abundance (*A*) from the pre-HAB period (2004–2020) to the post-HAB period (2021) with the following formulas (Iwasaki *et al*., 2016; Noda *et al*., 2017; Iwasaki & Noda, 2017; Ishida *et al*., 2023):

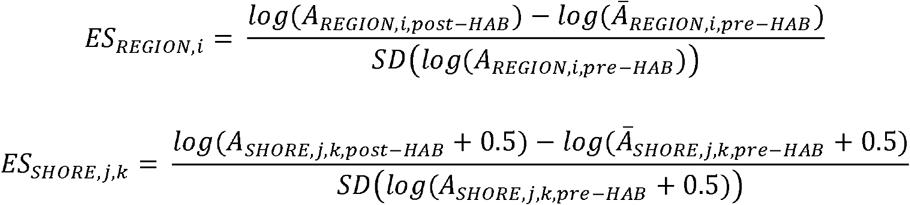

where *log* (*A*_*REGION,i,post − HAB*_) represents the log_10_(abundance) of each species in the post-HAB year, *log* (*Ā*_*REGION,i,pre−HAB*_) and *SD*(*log*(*A*_*REGION,i,pre−HAB*_)) represent the mean and standard deviation of the log_10_(abundance) of each species during the pre-HAB period, respectively; *log* (*A*_*REGION,j,k,post − HAB*_ + 0.5) represents the log_10_(abundance + 0.5) of each functional group at each shore in the post-HAB year, *log* (*Ā*_*REGION,j,k,pre − HAB*_ + 0.5) and *SD log* (*A*_*REGION,j,k,pre − HAB*_ + 0.5) represent the mean and standard deviation of log_10_(abundance + 0.5) of each functional group at each shore during the pre-HAB period, respectively. An effect size greater than 1.96 in absolute value suggests that the abundance in the post-HAB year changed significantly compared with the pre-HAB period.

These effect size approaches provide an accurate assessment of the immediate impact of the HAB because the effects of environmental stochasticity other than HAB are reduced in two ways: 1) the reference state abundance (abundance when not affected by the HAB) in this study is determined as a multi-year average, so the effect of environmental stochasticity other than the HAB on the change in abundance before and after HAB is small, and 2) the effect of HAB on abundance is assessed as the relative magnitude of the change in abundance before and after the HAB to the magnitude of the temporal variation of abundance in a normal year.

To evaluate whether spatial variations of the HAB impacts are related to the spatial distribution of cell density of HAB species, we employed scatter diagrams and a linear modelling to detect the relationship between *ES*_*HORE*_ and cell density of HAB species. For each functional group, we used *ES*_*HORE*_ as the dependent variable, and the cell density of *Karenia* spp. at each shore as the independent variable. All statistical analyses were conducted in R v. 4.3.1 (R Core Team, 2013).

## 3 Results

### 3.1 Effect of HAB on population sizes of different species and functional groups

Overall, the effect size at the regional scale had both positive and negative values (Fig. 3). Associations between HAB and macroalgae varied by species, and the effect size of *G. furcata* and *P. yendoi* were less than –1.96 (Fig. 3). However, the effect size of three species of molluscan grazers were all negative and less than –1.96 (Fig. 3). In addition, the effect sizes of *Chthamalus dalli* and *Nucella lima* indicated they were not significantly affected by the HAB.

**Fig. 3.**
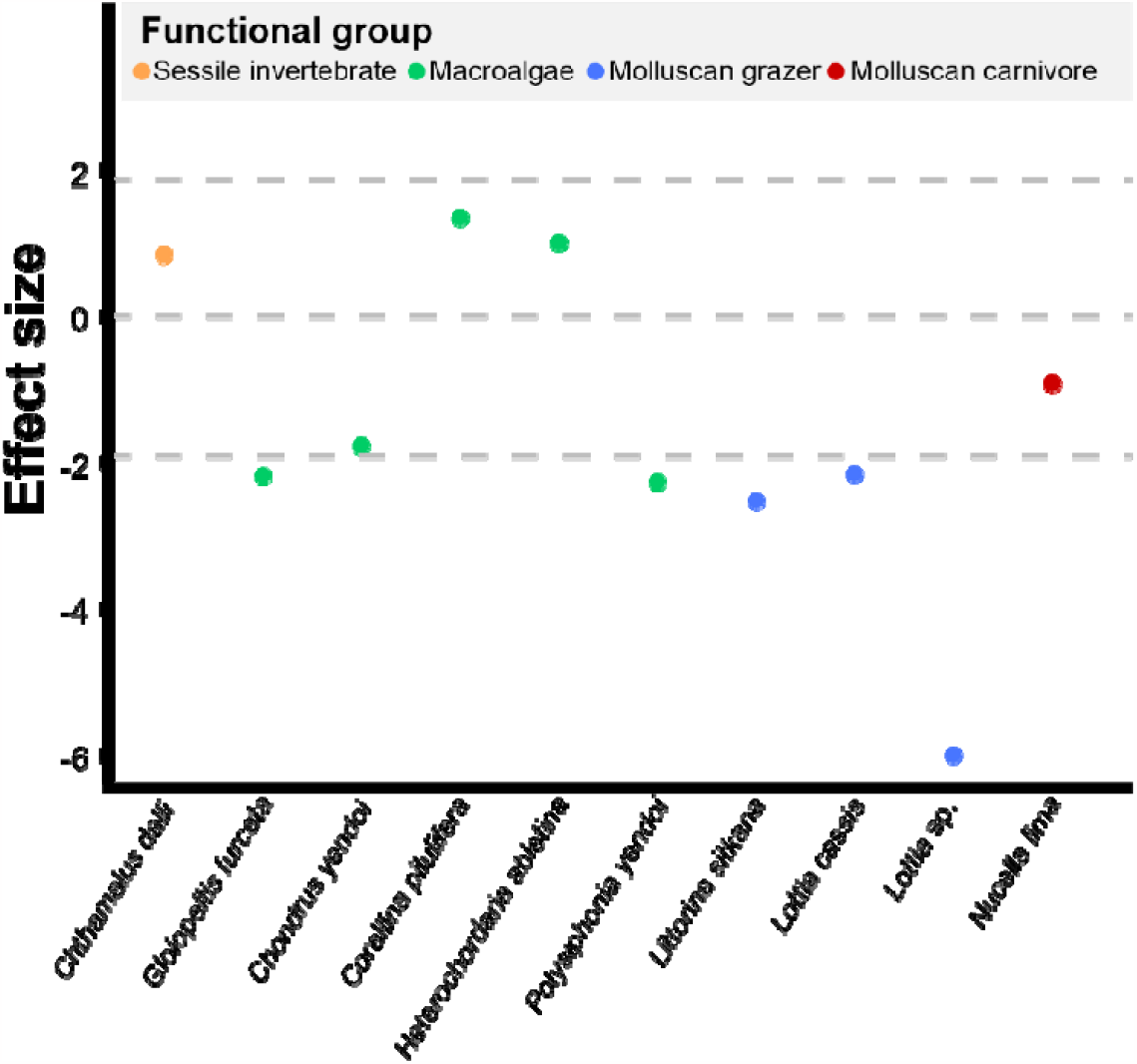
(color should be used in print). The effect size of different species at the regional scale. The dashed gray lines from top to bottom represent 1.96, 0, and –1.96, respectively. An effect size of greater than 1.96 in absolute value indicates a statistically significant difference in the population size before and after the HAB.

In terms of functional groups at the shore scale, the effect size of sessile animals and macroalgae had both positive and negative values among the shores, but the absolute value of the effect size of these two functional groups were less than 1.96, indicating that they were not significantly affected by the HAB at the shore scale (Fig. 4). However, the effect sizes of molluscan grazers were all negative, and the effect size was significantly negative at three shores (Aikappu, Mabiro, and Nikomanai) (Fig. 4). In addition, the effect size of molluscan grazers at Nikomanai was obviously less than the effect size of molluscan grazers at the other shores.

**Fig. 4.**
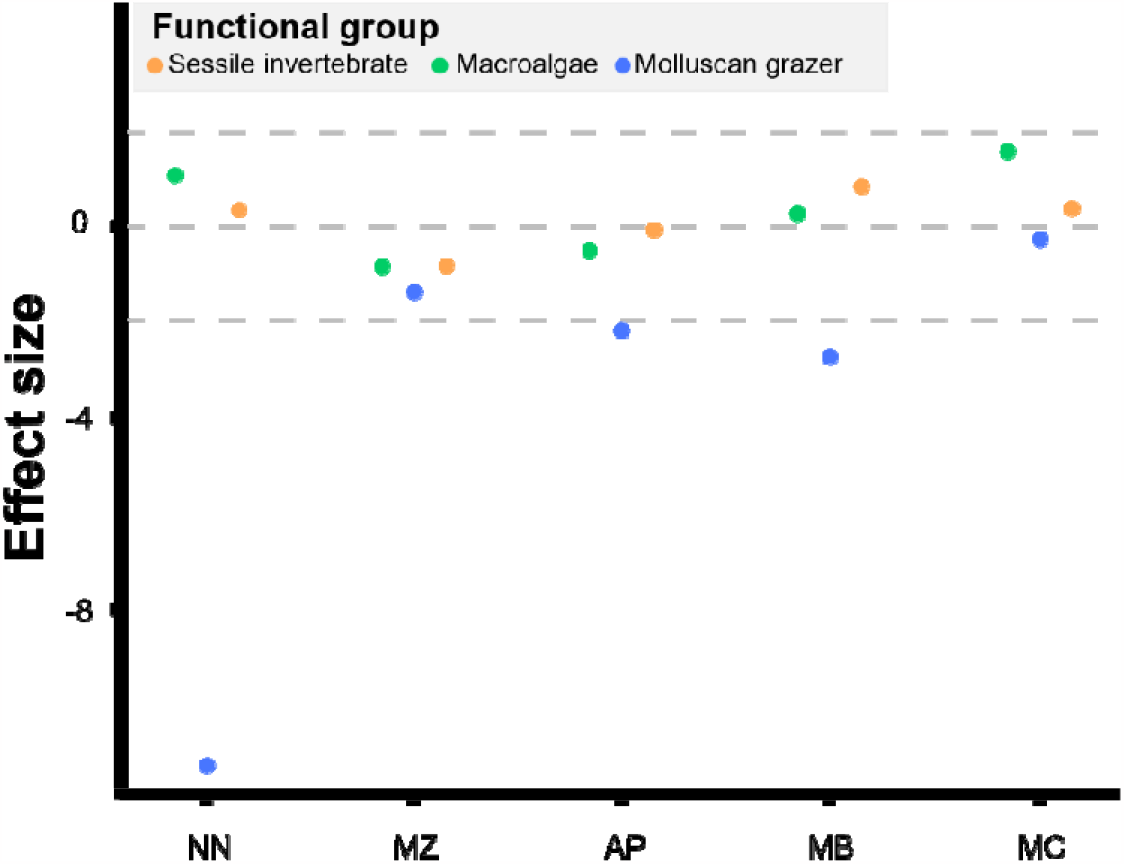
(color should be used in print). The effect size of different functional groups at the shore scale. The dashed gray lines from top to bottom represent 1.96, 0, and –1.96, respectively. An effect size of greater than 1.96 in absolute value indicates a statistically significant difference in the population size before and after the HAB. AP, Aikappu; MB, Mabiro; MC, Mochirippu; MZ, Monshizu; NN, Nikomanai.

### 3.2 The relationship between effect size and cell density of *Karenia* spp. among functional groups

The response of effect size to cell density of *Karenia* spp. varied among functional groups (Fig. 5). Only the effect size of a molluscan grazer was significantly negatively correlated with cell density (*p* < 0.01, Fig. 5). In addition, macroalgae and sessile invertebrates showed a low sensitivity to cell density (Fig. 5), and no significant relationships between effect size and cell density of *Karenia spp* were found in the linear modeling. (*p* > 0.05, Fig. 5).

**Fig. 5.**
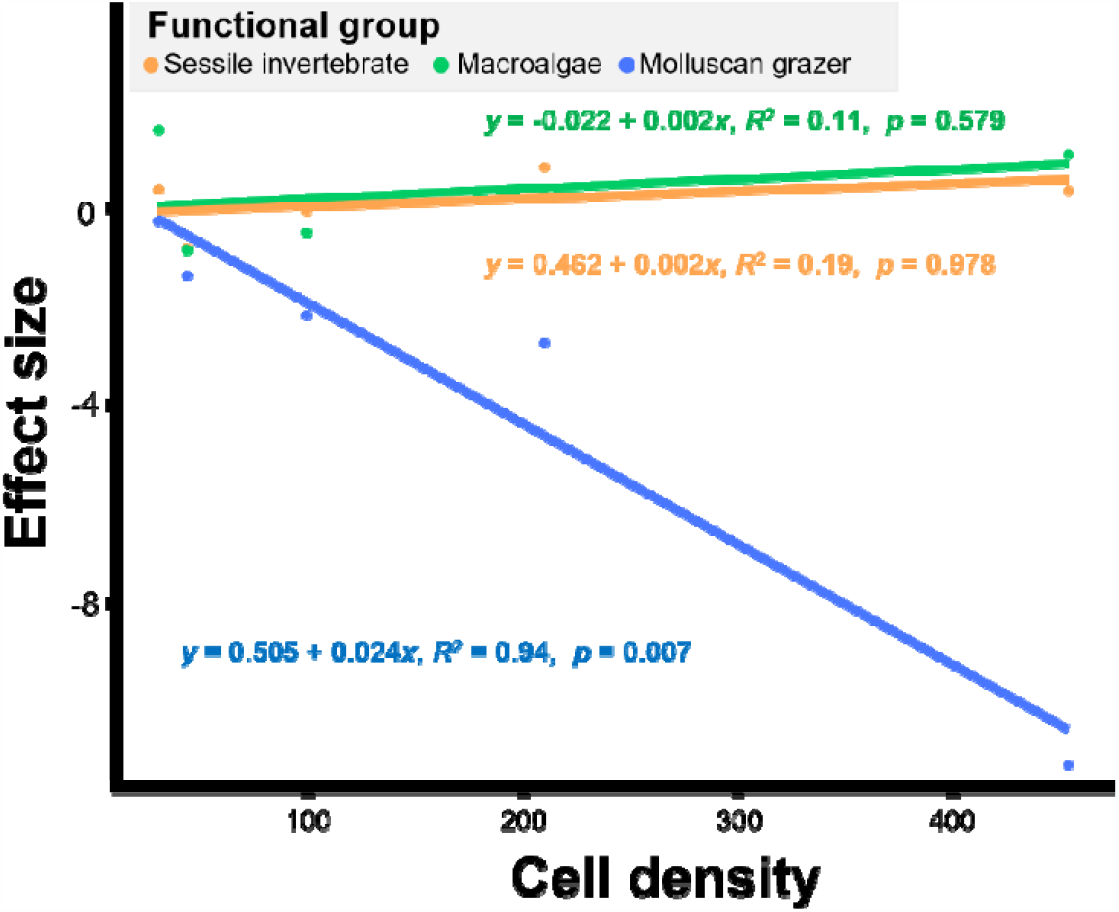
(color should be used in print). The relationship between of three functional groups (macroalgae, molluscan grazers, and sessile invertebrates) and the cell density of *Karenia* spp. at each shore. For the regression analyses, the cell density (cells/mL) of *Karenia* spp. at each shore and of each functional group were treated as independent and dependent variables, respectively.

## 4. Discussion

In this study, the immediate impact of the 2021 HAB in southeast Hokkaido on the population sizes of 10 common species and abundance of three functional groups in a rocky intertidal community was examined. Census data from 17 years before the HAB and within one month following the HAB were examined for 25 rocks at five shores along the coastline of southeast Hokkaido, Japan. Our results showed that two of five macroalgae species and all three molluscan grazer species significantly declined at the regional scale after the HAB occurred. In addition, only molluscan grazers declined significantly at three shores after the HAB. Furthermore, the effect size of molluscan grazers at the shore scale was significantly negatively correlated with cell density of *Karenia* spp.

While various year-to-year environmental fluctuations may drive community dynamics in rocky intertidal assemblages (Thompson *et al*., 2002; Menge *et al*., 2003; Ishida *et al*., 2023), their influences on the results of this study should be small. From October 2020 to October 2021, there was no other disturbance that would cause massive mortality (e.g., earthquake, tsunami, typhoon, ice scouring, marine heatwave, etc.) on the Pacific coast of southeast Hokkaido other than the HAB. Furthermore, other environmental factors that determine the population dynamics of rocky intertidal species, such as air temperature, sea water temperature, coastal topographic properties, and microtopography of rock surfaces were stable during that period. In addition, *ES*_*HORE*_ of other functional groups (macroalgae and sessile invertebrates) that were not affected by the HAB did not show any obvious consistent spatial patterns or significant variation, which was another evidence that the effects of environmental fluctuations on community dynamics were week.

Three molluscan grazers (*Littorina sitkana, Lottia cassis*, and *Lottia* sp.) and two macroalgae (*Gloiopeltis furcata* and *Polysiphonia yendoi*) significantly declined in abundance at the regional scale, but other macroalgae species and the sessile invertebrate and molluscan carnivore were unaffected. This difference can presumably be explained by differences in species-specific tolerance to biological toxins produced by *K. selliformis*, which is generally thought to produce gymnodimines and cause mass mortalities in a wide range of marine life (Ji *et al*., 2020), although there currently is no unified explanation for the mechanism of the lethal effect of *K. selliformis* on a wide range of marine organisms. A non-gymnodimine-producing phylotype of *K. selliformis* that causes massive fauna mortality by producing large amounts of long-chain polyunsaturated fatty acids and/or as-yet uncharacterized highly toxic compounds has also been reported (Mardones *et al*., 2020). Previous studies demonstrated that damage from *K. selliformis* differs among taxonomic groups. For example, *K. selliformis* has been shown to have a lethal effect on many mollusks, including octopus, limpets, and snails (Vellojin *et al*., 2023), as well as lethal effects on two juvenile kelp sporophytes *Saccharina japonica* and *S. sculpera* (Natsuike *et al*., 2023). However, there are no reported negative effects of *K. selliformis* on barnacles. In addition, massive kills of fish (Arzul *et al*., 1995; Heil *et al*., 2001) and shellfish (MacKenzie, 1996; Hasegawa *et al*., 2022) caused by *K. selliformis* have been reported around the world, but some bivalves that feed on *K. selliformis* can survive when it is present, and gymnodimines can be found accumulated in their tissues (MacKenzie *et al*., 1996; Biré *et al*., 2002). Furthermore, there are no reported negative effects of *K. selliformis* on macroalgae. The reason for these differences may be that different species have different tolerances for the biological toxins produced by HAB species.

Many studies of the toxicity of other HAB-causing species also showed that fish, shellfish, and invertebrates have different tolerances to species, such as *Karenia brevis* (Landsberg *et al*., 2009) and *Karenia mikimotoi* (Li *et al*., 2019). Compared to the impacts of the HAB along the west coast of South Africa (Branch *et al*., 2013) and other disturbances with similar geographical scale such as oil spills (Southward and Southward, 1978; Le Hir and Hily, 2002; Peterson *et al*., 2003), earthquakes (Iwasaki *et al*., 2016), nuclear disaster (Horiguchi *et al*., 2016), and ice scouring (Petzold *et al*., 2014) that have caused massive mortality on most species in rocky intertidal communities, the HAB in southeast Hokkaido had more obvious species-specific impacts that reduced the abundance of molluscan grazers and some macroalgae species. In addition, fisheries damage from the HAB was high for sea urchins, but oysters, scallops, and crabs were less affected (Ji *et al*., 2020; Hasegawa *et al*., 2022). These observations indicate that such strong species-specific impacts are a unique feature of the consequences of the 2021 Hokkaido HAB dominated by *K. selliformis*.

The significant decline of *G. furcata* and *P. yendoi* at the regional scale did not manifest at the functional group level at the shore scale. The response of macroalgae to the HAB was not uniform at the species level, and some other species that were not included in the analysis at the species level on the regional scale may increase after the HAB occurred (Fig. 3), which may offset the decline of *G. furcata* and *P. yendoi* on the functional group level at shore scale caused by HAB. In addition, scale transition theory provides a systematic framework to explain the problem of scaling up local-scale interactions to regional-scale dynamics with field data (Chesson *et al*., 2005; Chesson, 2012). When evaluating the population dynamics of the same species at different spatial scales, the predictions of dynamics at different scales diverge due to the interaction between nonlinearity in population dynamics at the local scale and the spatial heterogeneity in abundance and environmental factors that influence population dynamics (i.e., the intensity of the HAB in this study) at the regional scale (Melbourne & Chesson, 2006; Chesson, 2012). Therefore, the inconsistency of the significance of the HAB on effect size at different scales may be a result of the uneven distribution of the abundance of these two species at the regional scale and the spatial heterogeneity of environmental factors that influence population dynamics. Consequently, the increase of other macroalgae species and scale transition theory are not mutually exclusive and may operate simultaneously.

The spatial variation of the HAB impact in this study can be explained by the spatial pattern of *K. selliformis* cell density in offshore waters. Similarly, for an HAB along the coast of South Africa, the mortalities of several intertidal species were higher where the HAB intensity was stronger (Branch *et al*., 2013). In general, the severity of the impact from a disturbance (e.g., the resulting degree of population decline) is likely to depend primarily on the intensity of the disturbance (i.e., strength of forcing) (Iwasaki and Noda, 2018). Indeed, severity–intensity relationships within a single disturbance event have been documented in rocky intertidal organisms for various disturbance events occurring at scales from several to several tens of kilometers. For oil spills, the mortality of rocky intertidal organisms varies locally depending on levels of oil pollution (Southward and Southward, 1978; Le Hir & Hily, 2002; Peterson *et al*., 2003). For the Fukushima nuclear disaster, the abundance of surviving organisms decreased significantly with decreasing distance from the nuclear power plant (Horiguchi *et al*., 2016). For ice scarring, habitats that freeze for longer periods of time lose more biomass of sessile species in rocky intertidal habitat (Petzold *et al*., 2014).

The spatial variation of the impact of the HAB on the abundance of molluscan grazers in this study can be explained by the spatial pattern of *K. selliformis* cell density in offshore waters. Although the toxicological mechanisms of *K. selliformis* have not yet been clarified, this association between cell density and decrease of functional group abundance indicated that a high concentration of toxins in the HAB caused local mass mortality of molluscan grazers. For other dinoflagellates, previous studies indicated that the cell density of toxin-producing algae is always associated with the survival, mortality, or growth rate of molluscan species through various toxicological mechanisms (Yan *et al*., 2022). For example, the survival rate of the abalone *Haliotis discus* significantly decreased with increasing cell density of *Alexandrium pacificum* under a cultured condition (Zhang *et al*., 2018). This decline in survival rate may be caused by various toxicological mechanisms documented in bivalves, such as increased oxygen consumption, increased ammonia and phosphate excretion, and decreased Na^+^-K^+^ ATPase activity (Yan *et al*., 2022).

After the HAB in 2021, the abundance of molluscan grazers suddenly decreased at the shore scale, while the abundance of macroalgae and sessile animals remained unchanged. The loss of molluscan grazers could subsequently affect community dynamics through the trophic cascade (Lubchenco and Gaines, 1981; Menge, 2000). The decline in abundance of molluscan grazers will result in an increase in their food resource—macroalgae—and further decreases in the abundance of competitors of macroalgae, i.e., sessile invertebrates (e.g., barnacles) (Martins *et al*., 2008). Indeed, the experimental removal of molluscan grazers often results in increased macroalgae and decreased sessile invertebrates in rocky intertidal habitats (Moreno and Jaramillo, 1983; Phillips and Hutchison, 2008; Whalen *et al*., 2016).

Among the three molluscan grazers that suffered heavy HAB-induced population decline, *L. cassis* and *Lottia* sp. have a planktonic larval stage, whereas *L. sitkana* lacks a planktonic larval stage. This difference in life history may cause differences among species in the speed of population recovery at these three shores (Sahara *et al*., 2016). For species capable of larval dispersal, population recovery at Aikappu, Mabiro, and Nikomanai should benefit from larvae supplied from undamaged populations near each shore. On the other hand, the population recovery of non-planktonic species would completely depend on local reproduction. Thus, the speed of recovery at shores where molluscan grazers have significantly declined may be faster for populations of *L. cassis* and *Lottia* sp. than for *L. sitkana*.

## 5. Conclusion

In this study, long-term census data from five shores were used to evaluate the impacts of the 2021 HAB in southeast Hokkaido on the rocky intertidal community. Only two of five macroalgae species, *G. furcata* and *P. yendoi*, and three molluscan grazer species, *L. sitkana, L. cassis*, and *Lottia* sp., had significant population declines at the regional scale. At the functional group level at the shore scale, molluscan grazers declined significantly at three of five shores. In addition, the effect size of molluscan grazers was significantly negatively correlated with cell density of *Karenia* spp. These results imply that the effects of this HAB were highly species-specific and related to the density of the HAB-causing species in offshore waters. Because the toxicity of *K. selliformis* is still unclear, the mechanisms of species-specific and spatial variations of the HAB effects cannot be explained.

After the sudden loss of grazer species, increasing coverage of macroalgae and decreasing coverage of sessile animals resulting from the trophic cascade are predicted. Because of differences in larval dispersal ability, population recovery is expected to be faster in grazer species with planktotrophic larvae than in species with direct development.

## Acknowledgements

We are grateful to all the researchers and students who helped with field data collection, without which our study would not have been possible. This study received generous support and encouragement from local fishermen and the fisheries office of the Fisherman’s Cooperative Association in Akkeshi.

## Author contributions

The study was conceived by YY and TN. The data were collected by YY, TN, and other members of the long-term census project. The statistical analyses were performed by YY. The figures were prepared by YY. YY and TN wrote the paper.

